# A hierarchical abscission program regulates reproductive allocation in *Prunus yedoensis* and *Prunus sargentii*

**DOI:** 10.1101/2025.07.08.663657

**Authors:** Woo-Taek Jeon, Jeong-A Kim, Ahyeon Cheon, Shawn S. Y. Lee, Joohyun Kang, Jung-Min Lee, Yuree Lee

## Abstract

- organ abscission is essential for optimal reproduction, yet its regulation in perennial woody plant species is poorly understood. To investigate how abscission is spatially and temporally regulated during reproduction, we analyzed five sequential abscission events in the cherry species *Prunus yedoensis* and *Prunus sargentii*: abscission of the petals, calyces, flower pedicels, fruit pedicels, and peduncles.
- The abscission zone (AZ) of the calyx formed *de novo* upon activation, whereas other AZs were pre-formed but developmentally arrested. Localized ethylene responsiveness reactivated these zones, promoting cell division, differentiation of residuum and secession cells on either side of the AZ, and lignin deposition in some cases. This progression was accompanied by reactive oxygen species accumulation and pH shifts.
- We observed species-specific differences during early floral abscission: *P. yedoensis* shed petals rapidly in a pollination-independent manner, whereas *P. sargentii* retained petals on unpollinated flowers, which later abscised with the pedicel, potentially extending the fertilization window.
- Both species employed a post-fertilization checkpoint via fruit pedicel abscission to selectively eliminate small, slow-growing fruits. These findings reveal that *Prunus* species coordinate a hierarchical abscission program functioning as a multilayered reproductive filter, progressively refining investment decisions to determine the final fruit set.

## Introduction

Abscission is a biological program by which entire organs—such as leaves, flowers, and fruits—are actively separated from the plant body at defined cleavage sites known as abscission zones (AZs) (Addicott, 1982). This evolutionarily conserved mechanism serves diverse functions across kingdoms in processes including development, reproduction, and defense (Mierzwa and Gerlich, 2014, Golan and Pringle, 2017, Olsson and Butenko, 2018, Baban et al., 2022).

Abscission proceeds through a series of coordinated cellular events within the AZ: the differentiation of specialized cell layers, the acquisition of competence to perceive and respond to abscission signals, and the enzymatic remodeling of the cell wall to enable cell separation and organ detachment (Kim et al., 2019).

In addition to these established steps, recent studies in Arabidopsis (*Arabidopsis thaliana*) have revealed an additional layer of cellular differentiation within the AZ. Upon activation, cells in the AZ further specialize into two distinct types: secession cells (SECs), located on the side of the organ set to detach, and residuum cells (RECs), located on the plant body side and remaining after abscission (Lee et al., 2018). SECs form a lignin brace that spatially confines cell wall–degrading enzymes to the detachment interface, ensuring controlled separation. Disruption of this structure impairs proper organ shedding (Lee et al., 2018, Crick et al., 2022). Following organ detachment, RECs, which originally derived from mesophyll cells, acquire a new identity, transdifferentiating into epidermal-like cells that deposit a protective cuticle layer that functions as a barrier against water loss, mechanical damage, and pathogen invasion (Lee et al., 2018, Wen et al., 2025).

The activation of these cellular programs within the AZ is tightly regulated by upstream phytohormonal and signaling networks. Abscission is initiated through the coordination of intrinsic developmental cues and environmental stimuli, which converge to activate diverse signaling messengers in a spatiotemporally controlled manner. Among these factors, phytohormones such as ethylene and abscisic acid (ABA) act as positive regulators of abscission, whereas auxin and cytokinin function antagonistically to suppress abscission (Taylor and Whitelaw, 2001, Patharkar and Walker, 2018).

These phytohormonal cues ultimately feed into a core genetic program that executes abscission at the cellular level. A key component of this program is the INFLORESCENCE DEFICIENT IN ABSCISSION (IDA) pathway, in which the IDA mature peptide binds to the receptor-like kinases HAESA (HAE) and HAESA-LIKE2 (HSL2) (Butenko et al., 2003, Stenvik et al., 2008). This binding activates a mitogen-activated protein kinase (MAPK) cascade consisting of the MAPK kinases MKK4 and MKK5 that activate the MPKs MPK3 and MPK6, which promotes the expression of genes encoding cell wall–remodeling enzymes such as polygalacturonases and cellulases that are required for cell separation (Cho et al., 2008, Kumpf et al., 2013). Ethylene acts upstream of this module by inducing *IDA* expression, thereby linking phytohormonal signaling to the activation of the cell separation program (Butenko et al., 2006, Ying et al., 2016, Rai et al., 2021). Through this connection, the IDA– HAE/HSL2 module integrates developmental status with phytohormone responsiveness, forming a tightly controlled regulatory framework for abscission.

In addition to this phytohormone-driven transcriptional cascade, intracellular physiological signals, including reactive oxygen species (ROS) and cytosolic pH changes, modulate abscission by shaping the cellular microenvironment and influencing local enzymatic activity. ROS contribute to both the spatial and temporal regulation of abscission: spatially, they promote the localized polymerization of the lignin brace within SECs (Lee et al., 2018); temporally, the redox balance between superoxide and hydrogen peroxide, which is modulated by the secretory manganese superoxide dismutase MSD2, determines the onset of abscission (Lee et al., 2022). In parallel, the gradual cytosolic alkalinization of AZ cells is associated with both ethylene-dependent and ethylene-independent pathways, both of which are thought to regulate enzymatic activities or act as cues for gene expression (Sundaresan et al., 2015, He et al., 2023).

Given the tight coordination between abscission and developmental signaling, its functional significance extends beyond cellular execution to broader physiological contexts. In particular, abscission plays a critical role in plant reproduction, by removing floral organs that have completed their function after fertilization, as well as by enabling seed dispersal at later stages (Estornell et al., 2013). Much of our current molecular understanding of abscission comes from studies in Arabidopsis, which have identified key regulators and signaling pathways. However, its self-fertilizing nature results in relatively simple floral abscission dynamics that are largely uncoupled from fertilization outcomes, limiting its utility for investigating how abscission is regulated in response to reproductive success, such as whether fertilization or embryo development has occurred. In addition, its production of dry, silique-type fruits restricts its relevance for understanding abscission during the development or detachment of fleshy fruits. Although recent studies in crops such as rice (*Oryza sativa*) and tomato (*Solanum lycopersicum*) have expanded the molecular framework of abscission (Wu et al., 2023, Li and Su, 2024), the physiological roles of abscission in shaping reproductive efficiency remain underexplored, particularly in perennial or cross-pollinating species with more complex reproductive strategies.

To overcome these limitations, the genus *Prunus* offers a valuable system that combines reproductive complexity with broad biological relevance. It includes both economically important fruit crops and ornamental species, such as peaches (*Prunus persica*), apricots (*Prunus armeniaca*), almonds (*Prunus dulcis*), and cherries (*Prunus avium* and others) (Das et al., 2011, Dal Martello et al., 2023). Among these, ornamental cherries are particularly notable for their profuse flower production and strong reliance on cross-pollination, a combination that necessitates selective regulation over reproductive investment (Fairchild, 1911). Indeed, although profuse flowering increases the likelihood of successful fertilization, it may also result in more potential fruits than the plant is able to physiologically support. Limited nutrient availability and developmental capacity prevent all flowers from maturing into fruits. In this context, abscission likely serves as a developmental checkpoint, modulating reproductive output in response to both fertilization success and internal resource constraints. These features make the genus *Prunus* a compelling system in which to investigate how plants coordinate organ shedding with reproductive and physiological cues.

Despite this potential, relatively little is known about how abscission is regulated across developmental stages in *Prunus*, particularly during early phases of reproduction. Most existing studies in *Prunus* have focused on fruit maturation and detachment (Wittenbach, 1970, Wittenbach, 1975, Zanchin et al., 1995, Qiu et al., 2021). These studies showed that fruit detachment can occur either at the fruit–pedicel junction or at the pedicel–peduncle (PP) junction, and that the site of separation is developmentally regulated. Specifically, abscission tends to occur at the PP junction during early stages of fruit development, whereas it typically takes place at the fruit–pedicel junction in more mature fruits (Wittenbach, 1970). In *P. avium*, early activation of abscission at the PP junction varies among individuals and is correlated with higher rates of embryo abortion (Qiu et al., 2021), suggesting that abscission may act as a checkpoint to eliminate reproductively unsuccessful fruits. However, how earlier abscission events, particularly those involving floral organs, are developmentally coordinated with reproductive status remains poorly understood.

In this study, we investigated how *Prunus yedoensis* and *P. sargentii*, two ornamental cherry species, employ sequential and hierarchical abscission programs to regulate reproductive output. We show that *P. yedoensis* exhibits pollination-independent petal shedding, but subsequent abscission events of the calyx, flower pedicel, and fruit pedicel are selectively activated based on fertilization outcome and fruit development. In contrast, *P. sargentii* retains petals on unfertilized flowers and displays whole-flower abscission, revealing a distinct strategy for early reproductive filtering. Anatomical analysis further uncovered species-specific differences in lignin deposition and timing of AZ activation.

Together, our findings show that abscission operates as a layered developmental checkpoint that integrates developmental cues with reproductive outcomes to determine the fate of floral and fruit structures. This framework broadens the conceptual understanding of abscission beyond terminal detachment, offering new insights into its role as a key regulatory filter in reproductive optimization.

## Materials and methods

### Plant materials and growth conditions

Four-to five-year-old *P. yedoensis* trees used for pollination tests and ethylene treatments were cultivated in a pollinator-free greenhouse prior to anthesis to ensure controlled experimental conditions. The greenhouse was maintained at a temperature of 20–22 °C under natural light conditions. For the remaining experiments conducted under open-field conditions, specimens of *Prunus yedoensis* and *Prunus sargentii* var. *verecunda* aged seven years or older were collected from Seoul National University (37°27′36″N, 126°57′09″E) and Mt. Gwanak (37°26′44″N, 126°57′49″E), both located in the Gwanak district of Seoul, South Korea (Chang et al., 2004).

### Phenotypic assessment

To monitor abscission events in *P. yedoensis* and *P. sargentii* var. *verecunda* under open-field conditions, petal abscission, flower pedicel–peduncle abscission, calyx abscission, and fruit pedicel–peduncle abscission were scored daily at the branch and peduncle levels. The onset of each abscission event was recorded, and the number and proportion of each unit, including fully matured fruits, were quantified. Photographs corresponding to each abscission event were also collected using a KCS3-50 camera (Korea lab tech, gyeonggi-do, South Korea) and Leica M205FA stereomicroscope (Leica, Wetzlar, Germany). For seed imaging, the pericarp was manually removed from the fruit. Fruit and seed sizes were measured using ImageJ software (Abràmoff et al., 2004).

### Pollination treatment

For pollination treatment, *P. yedoensis* trees grown in a pollinator-free greenhouse were used. At 0–1 day post-anthesis (DPA), a subset of flowers corresponding to 0%, 25%, 50%, or 75% of the total number of flowers per tree, or all (100%) flowers were marked and subsequently hand-pollinated. Since *P. yedoensis* is self-incompatible, compatible pollen was obtained from five donor trees. Anthers were collected at 0 DPA, and pollen grains were released by inducing mechanical dehiscence of the anthers. The released pollen was then applied to the marked flowers.

### Scanning electron microscopy

Samples were fixed in 2.5% (v/v) glutaraldehyde in 0.1 M sodium phosphate buffer (pH 7.2), followed by vacuum infiltration for 1 h at room temperature. After the vacuum was released, samples were incubated overnight at room temperature. Subsequently, samples were washed three times in 1× phosphate-buffered saline (PBS) for 20 min each time and incubated in 1% (v/v) Triton X-100 at room temperature for 3 days. Post-incubation, samples were transferred to 1% (w/v) osmium tetroxide in 1× PBS and incubated overnight in the dark at 4°C. Samples were then washed three times in 1× PBS for 20 min each time and then subjected to ethanol dehydration using a graded series of ethanol solutions (30%, 50%, 60%, 70%, 80%, 90%, 95%, all v/v, and three times in 100%). Each step was conducted for a minimum of 2 h, with the final 100% ethanol step performed overnight. To maintain structural integrity, tissue specimens were dehydrated using a critical point dryer (EM CPD300, Leica, Vienna, Austria). The dried samples were affixed onto metal stubs with conductive carbon tape and coated with a thin platinum layer using a sputter coater (EM ACE200, Leica, Vienna, Austria) to improve electrical conductivity and imaging clarity. The prepared samples were then examined under high-vacuum conditions using a scanning electron microscope (JSM 6390LV, JEOL, Tokyo, Japan).

### ACC application via lanolin paste

1-Aminocyclopropane-1-carboxylate (ACC) was freshly dissolved in distilled water to a concentration of 1 mM, then incorporated into melted lanolin (pre-warmed to 60 °C) to yield a final concentration of 100 μM. A thin layer of the resulting lanolin paste was applied to specific floral organs, which were subsequently covered with plastic wrap to maintain contact: the calyx base for petal abscission treatment, and both the peduncle and pedicel sides for flower pedicel– peduncle abscission treatment. For control treatment, lanolin paste containing an equivalent volume of DMSO was applied to the corresponding sites.

### Resin sections for histological examination

Tissue samples were fixed overnight in 4% (v/v) glutaraldehyde (Sigma-Aldrich, Missouri, USA) prepared in 1× PBS (Bio-Rad Laboratories, California, USA). Following fixation, samples underwent dehydration through a graded ethanol series (30%, 40%, 50%, 60%, 70%, 80%, 90%, 95%, and 100%, all v/v). Infiltration with resin was carried out by gradually increasing the concentration of Technovit 7100 resin (Kulzer Technik, Hanau, Germany) in ethanol through a stepwise dilution series (1:5, 2:5, 3:5, 4:5, 5:5, 3.5:2.5, 4.5:2.5, 5.5:2.5, 6.5:2.5, 7.5:2.5, v/v), with each infiltration step conducted for 2 h. Fully infiltrated samples were embedded in resin, and 4-μm-thick sections were cut using a rotary microtome (Histocore AUTOCUT, Leica, Vienna, Austria). The sections were stained with 0.01% (w/v) toluidine blue (Thermo Fisher Scientific, Massachusetts, USA), and images were acquired using an Axioscope 5 microscope (Zeiss, Jena, Germany).

### DAB staining

Hydrogen peroxide accumulation was visualized by staining samples with 3,3′-diaminobenzidine (DAB; Sigma-Aldrich, Missouri, USA) (Straus et al., 2010). Vacuum infiltration was carried out using a 0.1% (w/v) DAB solution prepared in distilled water and adjusted to pH 5.8 with KOH. Samples were incubated in the dark for 6 hours to allow for DAB staining. After staining, tissues were fixed and chlorophylls were removed by incubating samples in a bleaching solution (ethanol:lactic acid:glycerol = 3:1:1, v/v/v) at 70 °C until complete decolorization was achieved. For petal AZs, samples were incubated in the bleaching solution at 70 °C for 1 hour. In the case of pedicel–peduncle and peduncle–branch AZs, the same 1-h incubation at 70 °C was performed four times with fresh bleaching solution exchanged after each step. Samples were mounted in bleaching solution and imaged using a Leica M205FA stereomicroscope (Leica, Wetzlar, Germany).

### Cytosolic pH visualization using BCECF

A 10-μM solution of BCECF-AM (Invitrogen, Massachusetts, USA), prepared from a 10-mM stock dissolved in DMSO (Sigma-Aldrich, Missouri, USA) and diluted in 1× PBS (Bio-Rad Laboratories, California, USA), was applied to the surface of tissue samples (Sundaresan et al., 2015). The samples were incubated in the dark for 30 min to allow dye uptake and then washed four times with 1× PBS to remove unincorporated dye. Fluorescence imaging was performed using an LSM 900 confocal microscope (Zeiss, Jena, Germany) with excitation at 488 nm and emission detection in the 505–525 nm range.

### Phloroglucinol–HCl staining for lignin detection

To visualize lignin accumulation, AZs from petals, pedicel–peduncles, and calyces were incubated in 3% (w/v) phloroglucinol in absolute ethanol at room temperature for 1 week, followed by incubation at 37 °C for an additional 3 weeks in a controlled chamber (Pomar et al., 2002). After incubation, 37% hydrochloric acid (HCl; Samchun Chemical Co., Ltd., Gyeonggi-do, South Korea) was applied to the tissues to initiate the staining reaction, with petals incubated for 1 min and pedicel–peduncle and calyx tissues incubated for 10 min. Samples were subsequently mounted in either 37% HCl or ethanol, and images were acquired using a Leica M205FA stereomicroscope (Leica, Wetzlar, Germany).

### Permeability assay using toluidine blue staining

To assess permeability, pedicel–peduncle and peduncle–branch RECs were immersed for 10 min in an aqueous solution containing 0.05% (w/v) toluidine blue O (Thermo Fisher Scientific, Massachusetts, USA) and 0.01% (v/v) Tween-20 (Sigma-Aldrich, Missouri, USA) (Tanaka et al., 2004). After staining, samples were rinsed twice with distilled water to remove excess dye. Images were acquired using a Leica M205FA stereomicroscope (Leica, Wetzlar, Germany).

### Petal color acquisition, correction, and visualization

To characterize petal coloration across developmental stages, from anthesis and petal abscission to flower pedicel–peduncle abscission, each sample was photographed using a KCS3-50 camera (Korea Lab Tech, Gyeonggi-do, South Korea) under standardized illumination within a matte black chamber. A color reference strip containing six predefined patches was included in each image for calibration. All image processing was conducted in MATLAB R2023b (MathWorks, Natick, MA, USA) using the Image Processing Toolbox. Reference regions were manually selected per image, and their red–green–blue (RGB) pixel values were converted into N×3 matrices to compute patch-wise mean values. A 3 × 3 linear correction matrix was then estimated via least-squares regression to map the measured RGB values to a predefined target set and applied to the entire image (Funt and Jiang, 2003). Corrected images were clipped to the 0–255 range, converted to 8-bit unsigned integers, and used for subsequent analysis.

To represent petal patterns in a compact and interpretable format, dominant colors were extracted from each image based on pixel frequency. Each image was smoothed using a Gaussian filter (σ = 1) to reduce noise. Thirty polygonal regions of interest (ROIs) were manually delineated per image to encompass intact petal areas while excluding regions in shadow or damaged regions. Binary masks were generated to isolate relevant pixels in each ROI. Extracted RGB values were converted to hue–saturation–value (HSV) color space, and pixels with V < 0.5 were excluded to minimize background interference. The frequency of the remaining RGB triplet values was computed, and the top 20 most frequent colors were selected for each ROI. From these, a representative set of 400 dominant RGB values was compiled across the 30 ROIs. Using the conversion values for RGB to HSV, all colors were sorted by H to enhance perceptual coherence (Chang and Mukai, 2022). The resulting sorted colors were visualized as a horizontal color bar placed beneath each corresponding petal image, enabling intuitive stage-wise comparison of observed petal color patterns (Zheng et al., 2022). MATLAB scripts used in this analysis are available upon request.

### Statistical analysis

All statistical analyses were carried out using GraphPad Prism software (version 10.5.0; GraphPad Software, https://www.graphpad.com/). Details regarding the number of technical and biological replicates and the specific statistical methods applied are included in the respective figure legends.

## Results

### Temporal and developmental coordination of sequential floral abscission in *P. yedoensis* and *P. sargentii*

To investigate how abscission is coordinated with reproductive development in *Prunus*, we monitored the timing and pattern of abscission events in *P. yedoensis* and *P. sargentii*, two ornamental cherry species. In these species, each inflorescence consists of 2–4 flowers, each borne on a separate pedicel attached to a common peduncle (Fig. 1a, b). This clear structural organization comprising floral organs, pedicels, and peduncles as separable units enabled precise staging of abscission events under open-field conditions.

**Fig. 1.**
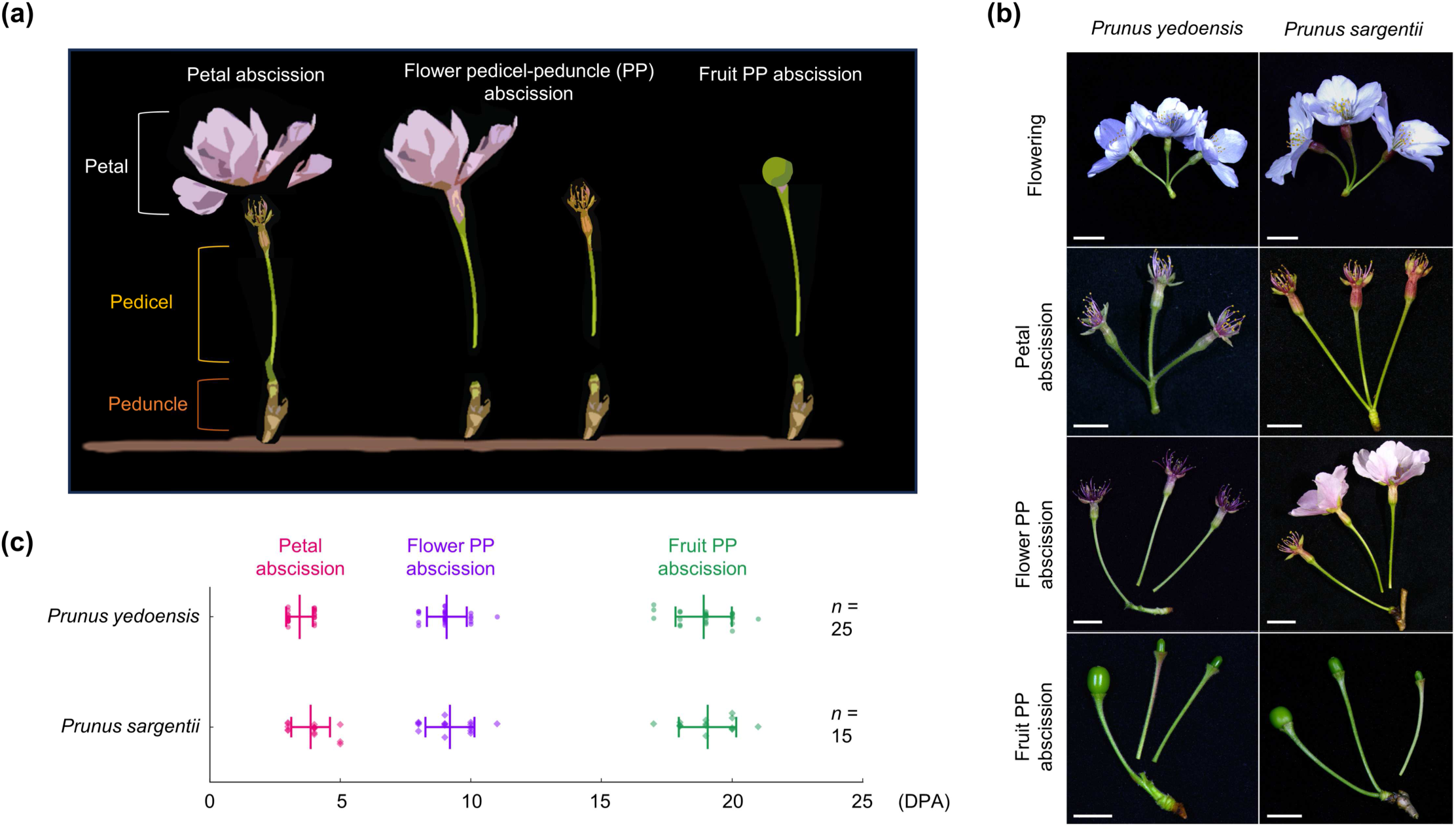
Phenotypic characterization of three distinct abscission types in *Prunus yedoensis* and *Prunus sargentii*. (a) Diagram depicting three types of abscission events in two *Prunus* species: petal abscission, flower pedicel–peduncle (PP) abscission, and fruit PP abscission. (b) Representative images of flowering and each abscission type. Scale bars, 1 cm. (c) Timing of the three abscission types in in two *Prunus* species. Data are presented as mean ± standard deviation (SD), with individual data points shown as distinct symbols. Quantification was performed under open-field conditions: for *Prunus yedoensis*, *n* = 25 branches from five independent trees were analyzed; for *Prunus sargentii*, *n* = 15 branches from three independent trees were analyzed. Data collected in 2025.

Both species exhibited a conserved sequence of abscission events: petal abscission around 3 days post-anthesis (DPA), followed by fruit pedicel–peduncle (PP) abscission at about 19 DPA (Fig. 1c). The latter abscission type involved detachment of the fruit-bearing pedicel from the peduncle, typically after the fruit had developed to a certain stage. Between these two events, we identified a third, intermediate abscission event, hereafter referred to as *flower PP abscission*, in which the pedicel of an unfertilized or aborted flower detaches from the peduncle (Fig. 1b). This event was initiated around 9 DPA at the same anatomical junction as fruit PP abscission but was developmentally distinct, as it involved a non-fruiting flower.

Although the timing of flower PP abscission was similar between the two species, its developmental context differed. In *P. sargentii*, flower pedicel detachment frequently occurred while the petals were still attached, whereas it typically followed petal shedding in *P. yedoensis* (Fig. 1b, c). These species-specific differences suggest that, although both species undergo three major abscission events (petal, flower PP, and fruit PP), their developmental coordination and regulatory thresholds are distinct.

### Petal abscission in *P. yedoensis* is developmentally timed and independent of pollination

To investigate the mechanisms underlying this species-specific coordination, we first focused on *P. yedoensis*. A striking feature of this species is its rapid, synchronous petal shedding shortly after anthesis. Given that floral structures primarily function to facilitate pollination (Real, 2012, Yuan et al., 2013), and that *Prunus* species depend on insect-mediated cross-pollination (Calzoni and Speranza, 1998, Yamane and Tao, 2009, Hansted et al., 2012, Dar et al., 2018), this pattern raised a key question: does the rapid and synchronized petal abscission seen in *P. yedoensis* reflect high pollination efficiency?

To address this question, we hand-pollinated flowers at defined levels (0%, 25%, 50%, 75%, or 100% of all flowers per tree) and monitored petal abscission under greenhouse conditions. Contrary to the common assumption that petal retention is linked to pollination success, flowers of *P. yedoensis* uniformly shed their petals even in the complete absence of pollination (Fig. 2a), suggesting that petal abscission is developmentally programmed. Pollinated and nonpollinated flowers had identical petal shedding kinetics (Fig. S1a, b), confirming that petal abscission is triggered independently of pollination and follows an intrinsic schedule initiated after anthesis.

**Fig. 2.**
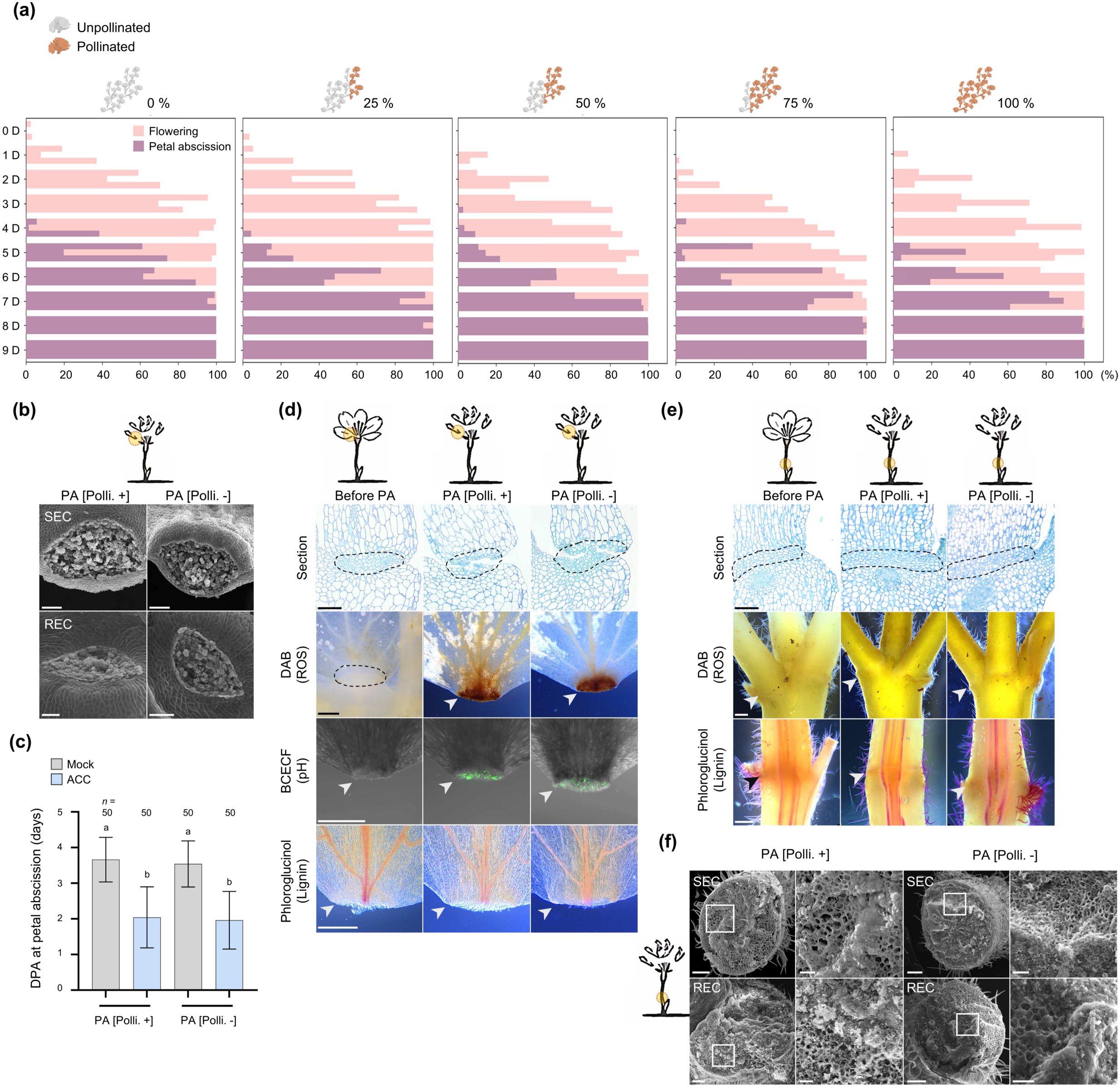
Petal abscission, but not pedicel–peduncle abscission, occurs independently of fertilization in *Prunus yedoensis*. (a) Temporal pattern of petal abscission in response to pollination. For each tree, the proportion of flowers that had undergone anthesis (representing flowering) was recorded, and cross-pollination was performed on 0%, 25%, 50%, 75%, or 100% of all flowers per tree at 0–1 day post-anthesis (DPA). The daily percentage of petal abscission events is also plotted for each pollination group. Each horizontal bar represents data from an individual tree (*n* = 3 trees). (b) Representative scanning electron micrographs showing residuum cells (RECs) and secession cells (SECs) comprising the petal abscission zone (AZ), following manual separation at the time of petal abscission (PA), with pollination (PA [Polli. +]) or without pollination (PA [Polli.−]) (*n* = 5 flowers). (c) Time to petal abscission following mock treatment or 1-aminocyclopropane-1-carboxylate (ACC) treatment. Pollinated and nonpollinated flowers were treated with either mock solution (DMSO) or 100 µM ACC. To account for the time required for fertilization, pollination was defined as 0 DPA, and treatments were applied at 1 DPA. Data are shown as means ± SD. Statistical analysis was performed using one-way analysis of variance (ANOVA) with Dunn’s post-hoc correction (*n* = 50 flowers). (d) Representative images of sections, DAB staining, BCECF staining, and phloroglucinol*–*HCl staining of petal AZs. Samples were collected at three stages: before petal abscission (Before PA), during petal abscission after pollination (PA [Polli. +]), and during petal abscission without pollination (PA [Polli. −]). Black dashed lines and arrowheads indicate the location of the AZ (*n* = 5 for section images; *n* = 21 for all other staining experiments). (e) Representative images of sections, DAB staining, and phloroglucinol*–*HCl staining of the flower pedicel– peduncle (PP) AZ. Samples were collected at three time points. Black dashed lines and arrowheads indicate the location of the AZ (*n* = 5 for section images; *n* = 21 for all other staining experiments). (f) Representative scanning electron micrographs showing RECs and SECs comprising the PP AZ, following manual separation during the PA [Polli. +] and PA [Polli. −]. The white squares indicate the regions shown at higher magnification at right (*n* = 5). Scale bars, 100 μm (b); 50 μm (d and e, higher magnification images in f); 500 μm (DAB, phloroglucinol*–*HCl, and BCECF images in d and e); 200 μm (main images in f).

Scanning electron microscopy (SEM) analysis supported this conclusion. Indeed, we observed clean separation across the AZ regardless of pollination status. Petals manually removed at the time of natural abscission showed uniformly smooth fracture surfaces in both SECs and RECs, with no sign of torn or collapsed tissue (Fig. 2b), indicating that cell separation proceeds cleanly across the AZ via programmed cell wall remodeling, independently of pollination. Application of ethylene, a key phytohormonal regulator of abscission (Brown, 1997, Meir et al., 2019), accelerated petal abscission (Figs. 2c, S2a), demonstrating that the petal AZ is developmentally primed and fully responsive to ethylene at this stage.

At the cellular level, Arabidopsis employs spatially localized ROS accumulation to regulate the timing and pattern of abscission (Lee et al., 2018, Lee et al., 2022). Similarly, in *P. yedoensis*, ROS accumulated in the AZ coinciding with the onset of abscission (Fig. 2d), suggesting a conserved mechanism. Beyond a marker of timing, ROS may trigger downstream events such as localized alkalinization, which modulates the activity of cell wall–remodeling enzymes (Sundaresan et al., 2015). Indeed, imaging with the pH-sensitive fluorescent dye BCECF (2′,7′-bis(2-carboxyethyl)-5-(and-6)-carboxyfluorescein) revealed a marked increase in fluorescence in the AZ during abscission (Fig. 2d), indicating localized alkalinization. Together, these findings suggest that ROS and pH act in concert to regulate the spatial and temporal progression of abscission. Unlike Arabidopsis, which uses lignin deposition to demarcate the AZ boundary (Lee et al., 2018), *P. yedoensis* exhibited no lignin accumulation in the petal AZ (Fig. 2d). This finding suggests that pH-mediated enzyme modulation, rather than lignin-based compartmentalization, contributes to the spatial control of abscission in this species.

In contrast, the PP AZ remained inactive at this stage. Anatomical analysis revealed that the PP AZ was already structurally differentiated when petal abscission occurred (Fig. 2e). However, SEM analysis of manually detached pedicels showed torn and irregular fracture surfaces (Fig. 2f), indicating that abscission had not yet been activated. Application of the ethylene precursor ACC also failed to induce pedicel abscission (Fig. S2b), suggesting that the AZ was not yet ethylene responsive. In support of this idea, we detected no ROS accumulation or lignin deposition in the PP AZ at this stage (Fig. 2e). These findings highlight a clear developmental separation between different types of AZs: while the petal AZ is actively executing abscission, the anatomically formed PP AZ remains dormant until later stages, underscoring the stage-specific activation of abscission programs in *P. yedoensis*.

### The divergent abscission paths of the calyx and pedicel are shaped by fruit development in *P. yedoensis*

Pollination-independent petal abscission leads to the retention of many unfertilized flowers, prompting the question of how the plants eliminate these non-productive remnants. Because these flowers, which retain reproductive organs and pedicels, offer no reproductive advantage, we hypothesized that flower PP abscission serves as clearance mechanism (Fig. 3a). Supporting this idea, the fruits from flowers that underwent flower PP abscission were significantly smaller than those that remained attached (Fig. 3b), indicating that flower PP abscission preferentially removes low-output flowers.

**Fig. 3.**
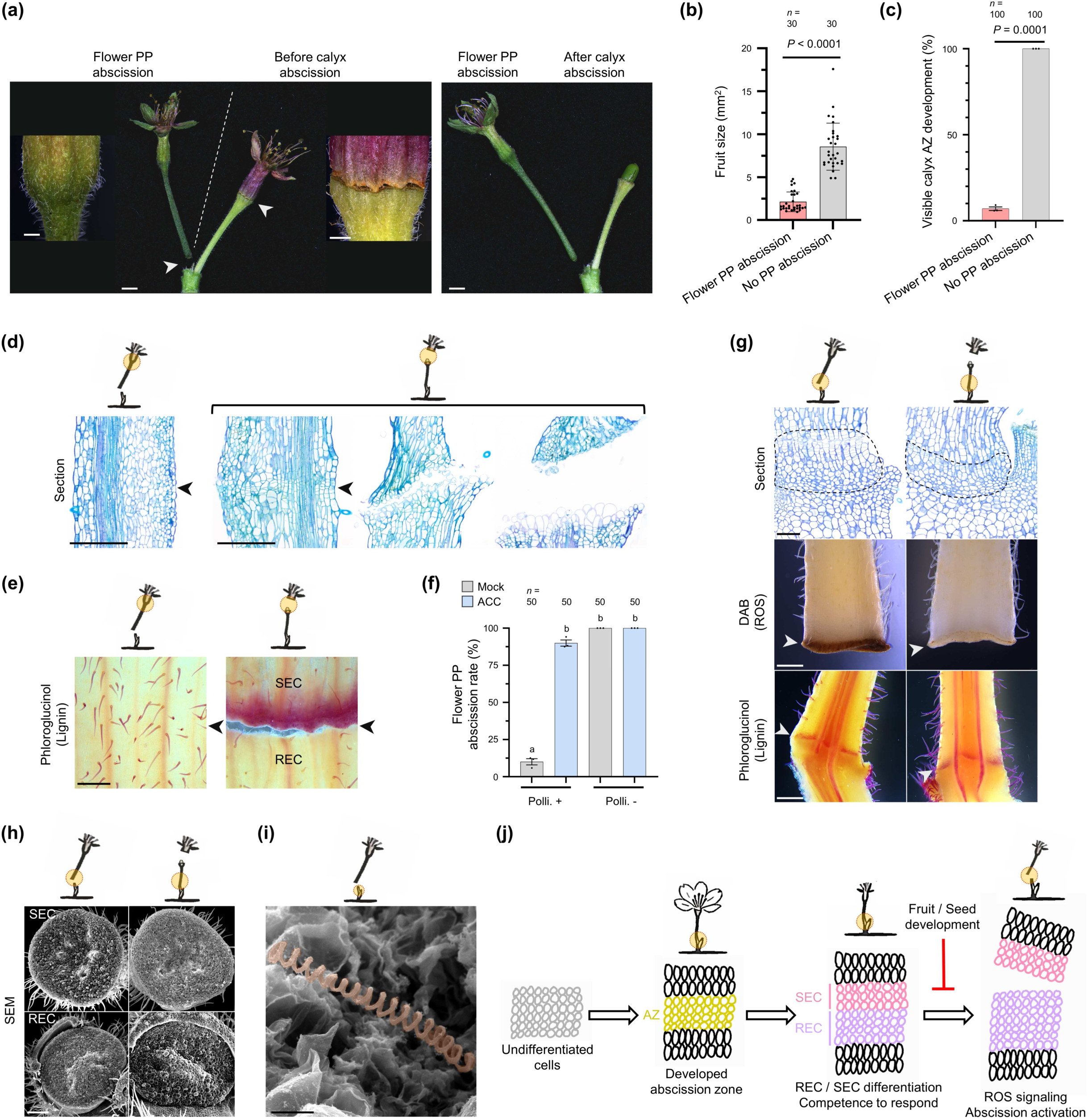
Reproductive success determines abscission type between flower pedicel–peduncle and calyx in *Prunus yedoensis*. (a) Representative images of flower pedicel–peduncle (PP) abscission and calyx abscission occurring on the same peduncle. Insets show higher-magnification views highlighting the calyx region in both abscission types. (b) Fruit size inside flowers at the time of flower PP abscission and inside flowers that did not undergo flower PP abscission. Sampling was conducted at 9– 11 DPA; flowers that did not undergo flower PP abscission subsequently proceeded to calyx abscission. Each dot represents an independent flower; data are shown as means ± SD. Statistical analysis was performed using the Mann–Whitney test (*n* = 30 flowers). (c) Proportion of flowers that developed a calyx abscission zone (AZ) as a function of whether flower PP abscission occurred. For flower PP abscission, observations were made at 9–11 DPA, when abscission is activated. In cases where PP abscission did not occur, samples were observed up to 20 DPA to track the development of the calyx AZ. Statistical analysis was performed using Welch’s *t*-test. Data are shown as means ± SD (*n* = 100 flowers). (d) Representative section images of the calyx AZ in flowers undergoing either flower PP abscission or calyx abscission. Black arrowheads indicate the expected location of the calyx AZ. (e) Representative phloroglucinol–HCl staining images for lignin detection in the calyx AZ. Staining was performed on calyx AZs from flowers undergoing either flower PP abscission or calyx abscission. Samples were collected at time points corresponding to the physiological activation of each AZ. Black arrowheads indicate the expected location of the calyx AZ (*n* = 12). (f) Flower PP abscission rate following mock treatment or ACC treatment. Pollinated and nonpollinated flowers were treated with either mock solution (DMSO) or 100 µM ACC. Treatments were applied at 5 DPA, after the completion of petal abscission but prior to the onset of flower PP abscission. The abscission rate was measured at 15 DPA, when flower PP abscission naturally occurs while fruit PP abscission has not yet been initiated. Statistical analysis was performed using one-way ANOVA with Dunn’s post-hoc correction (*n* = 50 flowers). (g) Representative images of resin sections, DAB staining, and phloroglucinol–HCl staining of the flower PP AZ from flowers undergoing either flower PP abscission or calyx abscission. Samples were collected at time points corresponding to the physiological activation of each AZ. Black dashed lines and white arrowheads indicate the location of the flower PP AZ (*n* = 5 for section images; *n* = 12 for all other staining experiments). (h) Representative scanning electron micrographs showing RECs and SECs comprising the PP AZ, following manual separation at the time of flower PP abscission or calyx abscission. Samples were collected at time points corresponding to the physiological activation of each AZ (*n* = 5). (i) Representative scanning electron micrographs of RECs from flower PP abscission, highlighting exposed spiral vessels in the sectioned tissue shown in orange pseudocolor. Samples were collected at time points corresponding to the physiological activation of the flower PP AZ (*n* = 5). (j) Diagram of the development and activation of the flower PP AZ. During anthesis, the AZ undergoes structural development. Around the time of petal abscission, REC and SEC differentiation occurs, together with lignin brace formation and acquisition of ethylene responsiveness (competency). Subsequently, reactive oxygen species (ROS) signaling is activated in the SECs, leading to the initiation of abscission. By contrast, when fruit and/or seed development proceeds successfully, this activation step is bypassed, and calyx abscission occurs instead. Scale bars, 10 mm (main images in a), 1 mm (higher-magnification images in a), 500 μm (d, e, DAB and phloroglucinol images in g), 50 μm (sections in g), 200 μm (h), 20 μm (i).

Notably, in flowers that remained attached, we observed an alternative abscission event at the calyx, the whorl of sepals enclosing the floral organs during early development (Figs. 3a, S3). This calyx abscission event was absent in flowers that had undergone flower PP abscission, which lacked any visible boundary between the calyx and pedicel (Fig. 3c). Longitudinal sections confirmed this observation, as we saw no anatomical trace of a calyx AZ in these flowers (Fig. 3d). In contrast to the PP AZ, which is pre-formed but developmentally dormant, the calyx AZ thus appears to be absent until later stages.

However, in flowers that escaped flower PP abscission and remained attached, a *de novo* AZ emerged at the base of the calyx during fruit development, comprising three to four layers of narrow cells (Fig. 3d). This zone displayed hallmark features of AZ activation: REC expansion, separation from SECs, and progressive REC enlargement. Moreover, lignin deposition occurred specifically in the SEC layer of the calyx AZ (Fig. 3e), echoing the situation in Arabidopsis, in which lignin functions as an apoplastic brace to spatially constrain cell wall degradation.

In contrast to the calyx AZ, which forms only after fertilization, the PP AZ is already anatomically established by anthesis but remains ethylene-insensitive at the time of petal shedding (Figs. 2e, S2b). Flower PP abscission occurred selectively among flowers on the same peduncle, indicating localized regulatory control. To clarify whether this selectivity arises from differences in ethylene competence, signaling input, or structural inhibition, we compared ethylene responsiveness between pollinated and nonpollinated flowers. Abscission was markedly less frequent in pollinated flowers but occurred frequently in nonpollinated ones (Fig. 3f). However, ACC treatment restored abscission in pollinated flowers to levels comparable to that seen in nonpollinated controls, ruling out structural constraints or differential competence. These findings suggest that selective PP abscission is governed primarily by signaling differences rather than anatomical or phytohormonal readiness.

At this stage, lignin began to accumulate in the SEC of the PP AZ, a feature absent from earlier stages, indicating structural maturation (Fig. 3g). The number of AZ cell layers also increased, especially within the REC region of abscising pedicels, suggesting enhanced tissue readiness. As in Arabidopsis (Lee et al., 2018), lignin deposition preceded organ separation and occurred irrespective of whether abscission ultimately took place. SEM analysis of manually separated pedicels further supported this notion, revealing smooth fracture surfaces and lignified xylem strands as the final physical link in both abscised and retained pedicels (Fig. 3h, i). These observations confirm that the PP AZ is structurally delineated and primed for separation, regardless of whether abscission is ultimately executed.

Notably, we observed a more pronounced increase in AZ cell layers in pedicels undergoing PP abscission, especially within the REC region, indicating enhanced maturation associated with activation (Fig. 3g). In addition, we detected ROS accumulation only in abscising pedicels (Fig. 3g), suggesting that ROS function as a terminal trigger for abscission. These findings suggest that the PP AZ progressively acquires structural readiness and ethylene responsiveness regardless of fertilization outcome, but undergoes further maturation upon seed abortion, including ROS activation, to initiate abscission (Fig. 3j).

Together, these results demonstrate that calyx and flower PP abscissions follow distinct temporal and regulatory trajectories. Calyx abscission is induced post-fertilization in coordination with fruit development, whereas pedicel abscission serves as a selective mechanism for the removal of non-productive flowers, integrating structural maturation, ethylene sensitivity, and ROS signaling.

### A post-fertilization abscission checkpoint controls fruit number in *P. yedoensis*

Flower PP abscission efficiently eliminates unfertilized flowers at early stages, but whether all fertilized flowers are maintained through fruit development remains unclear. Insect-mediated pollination is inherently variable. Although flower overproduction helps ensure reproductive success, excessive fruit retention under high pollination success could overwhelm resource availability and compromise reproductive efficiency.

We hypothesized that *P. yedoensis* employs a secondary filtering mechanism after fertilization to regulate final fruit load. To test this hypothesis, we tracked the fate of flowers and developing fruits across stages. Remarkably, a much of fruit PP abscission occurred after fruit development had already initiated, accounting for 41.9% of all flowers and 73.0% of calyx-abscised fruits (Figs. 1c, 4a). This post-fertilization abscission was strongly associated with smaller fruit size: abscised fruits were on average 38.5% as large as retained fruits (Fig. 4b), and their seeds were about 16.7% as large as those in retained fruits (Fig. 4c). These findings indicate that *P. yedoensis* employs post-fertilization fruit selection via fruit PP abscission to eliminate low-quality fruits, thereby optimizing resource allocation and enhancing reproductive efficiency. This filtering may reflect intrinsic fruit quality, inter-fruit competition, or both, acting as a checkpoint.

**Fig. 4.**
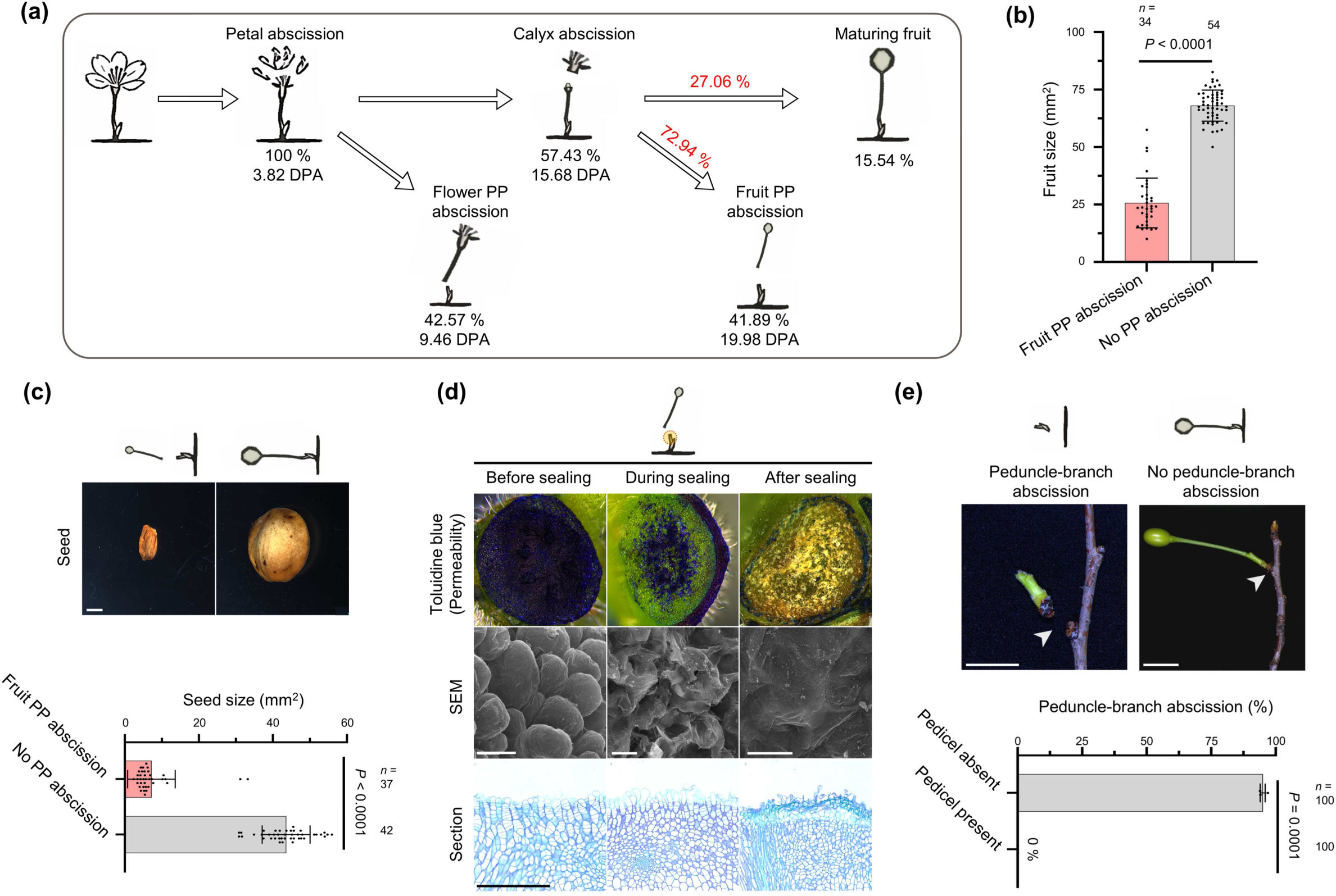
Reproductive success influences subsequent abscission types in *Prunus yedoensis*. (a) Diagram showing the timing of each abscission type and the proportion of flowers (or fruits) undergoing each abscission type. The percentages shown below each image represent the absolute proportion relative to the total number of flowers (or fruits), and the percentages shown above the arrows represent relative proportions. Data collected in 2025. (b) Fruit size at the time of fruit PP abscission in fruits that underwent fruit PP abscission and those that did not. Sampling was conducted at 19–22 DPA, corresponding to the period when fruit PP abscission occurs. Statistical analysis was performed using the Mann–Whitney test. Each dot represents an independent fruit; data are shown as means ± SD (*n* = 54 fruits). (c) Representative images of seeds (top) and seed size (bottom) at the time of fruit PP abscission in fruits that underwent fruit PP abscission and those that did not. Sampling was conducted at 19–22 DPA, corresponding to the period when fruit PP abscission occurs. Statistical analysis was performed using the Mann–Whitney test. Data are shown as means ± SD (*n* = 37 or 42 seeds each). (d) Representative images of toluidine blue staining, sections, and scanning electron micrographs illustrating the development of the protective layer in RECs of the PP AZ. Samples were collected at three stages: before sealing (at fruit PP abscission), during sealing (3 days after fruit PP abscission), and after sealing (6 days after fruit PP abscission). (*n* = 10 PP RECs for toluidine blue staining; *n* = 5 for sections and SEM analysis). (e) Representative images (top) and frequency (bottom) of peduncle–branch abscission. White arrowheads indicate the location of the flower peduncle–branch AZ. Statistical analysis was performed using Welch’s *t*-test (*n* = 100). Data are shown as means ± SD. Scale bars, 2 mm (c), 500 μm (toluidine blue staining and sections in d), 20 μm (SEM images in d), 1 cm (e).

Given the high frequency of post-fertilization abscission, maintaining tissue integrity at the exposed site is essential. To examine how *P. yedoensis* achieves this, we assessed surface permeability using toluidine blue staining. We observed a gradual decline in staining intensity over time, indicating that a protective barrier forms post-abscission (Fig. 4d). Unlike in Arabidopsis, where RECs transition into epidermal-like cells and develop a cuticle (Lee et al., 2018), *P. yedoensis* formed a yellowish, translucent material that spread over the abscised surface. SEM analysis revealed that this material originates from RECs and flows across cell boundaries (Fig. 4d). Longitudinal sections showed dynamic changes in the RECs, including outer cell expansion, inner cell divisions, and eventual collapse of the outermost layer. These observations suggest a multilayered, species-specific sealing mechanism involving coordinated cellular remodeling.

Strikingly, once all associated pedicels had abscised, the peduncle itself was also shed. Specifically, 95% of peduncles abscised when all pedicels were gone, whereas those retaining even a single pedicel remained intact (Fig. 4e). As with other abscission events, we detected evidence for ROS accumulation, as well as the formation of a protective layer on branch-side RECs (Fig. S4). Collectively, these findings reveal that *P. yedoensis* orchestrates a sequential and hierarchical abscission program, from the early removal of unfertilized flowers, to the post-fertilization elimination of underdeveloped fruits, to the eventual shedding of spent peduncles. This multilayered strategy ensures efficient reproductive investment and resource optimization throughout its reproductive phase.

### Species-specific adjustment of hierarchical abscission in *P. sargentii*

To determine whether the hierarchical sequence of abscission sequence observed in *P. yedoensis* represented a species-specific adaptation or a general reproductive strategy, we extended our analysis to the closely related species *P. sargentii*. When abscission events were tracked at the level of individual peduncles, *P. sargentii* displayed a similarly ordered progression: petal abscission occurred first, followed by flower PP abscission, calyx abscission, and finally fruit PP abscission (Fig. 5a). This stepwise filtering resulted in final fruit set accounting for less than 8% of the initial number of flowers (Fig. 5a). Notably, both species exhibited strikingly similar rates of fruit PP abscission among calyx-abscised fruits (*P. yedoensis*: 72.9%; *P. sargentii*: 73.2%), suggesting a conserved mechanism regulating final fruit load.

**Fig. 5.**
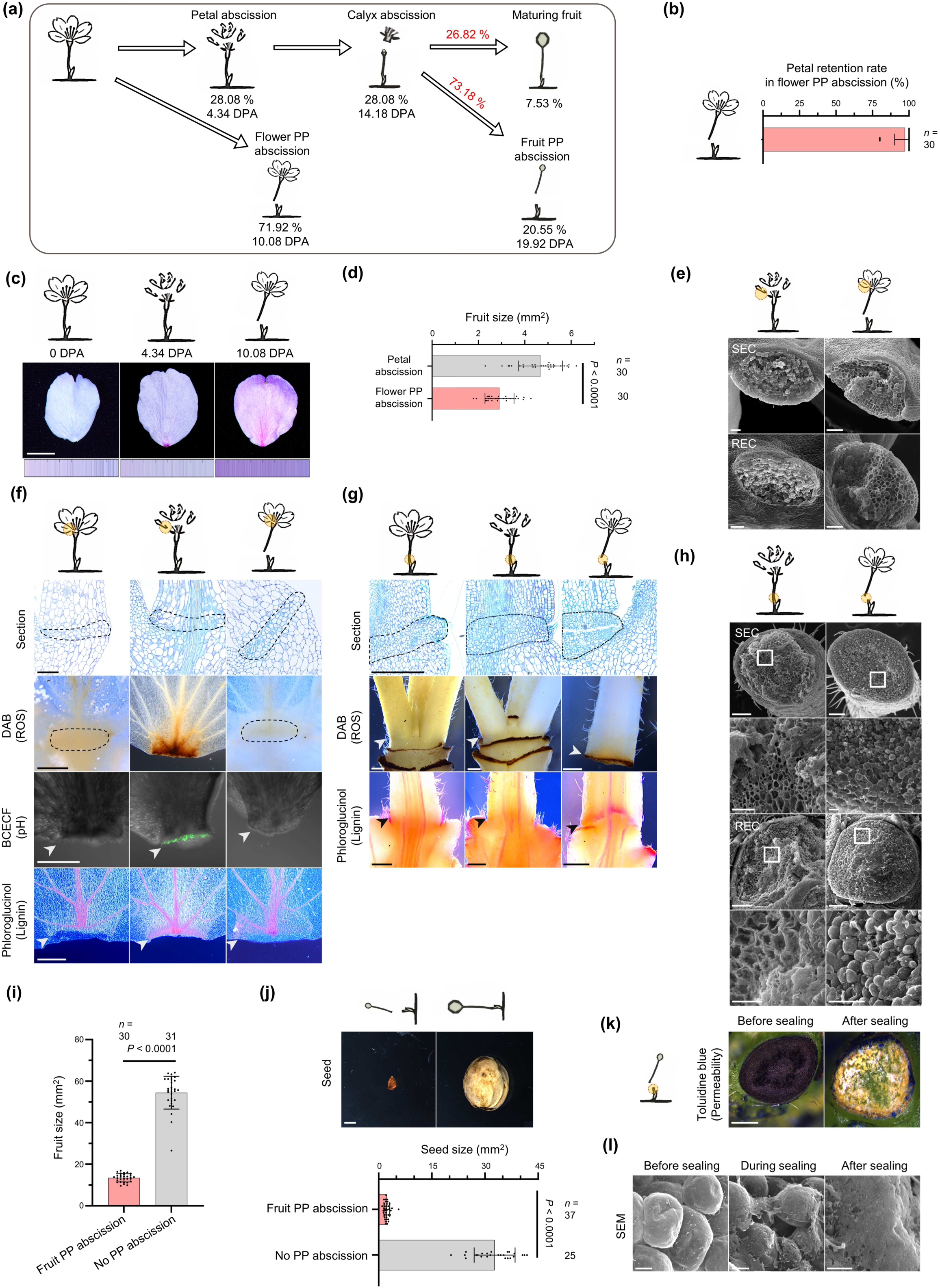
Choice between petal and flower pedicel abscission types in *Prunus sargentii* is guided by reproductive success. (a) Diagram showing the timing of each abscission type and the proportion of flowers (or fruits) undergoing each abscission type. The percentages shown below each image represent the absolute proportion relative to the total number of flowers (or fruits), and the percentages shown above the arrows represent relative proportions. Data collected in 2025. (b) Number of petals remaining on the flower at the time of flower PP abscission. Each dot represents an independent flowers; data are shown as means ± SD (*n* = 30 flowers). (c) Transition of petal color from anthesis to petal abscission and flower PP abscission. Representative images of petals are shown, and the color bars below each image depict the palette based on 400 measurements (*n* = 20 flowers). (d) Fruit size inside flowers at the time of flower PP abscission for flowers undergoing petal abscission and those undergoing flower PP abscission. Each dot represents an independent fruit; data are shown as means ± SD. Statistical analysis was performed using the Welch’s *t*-test (*n* = 30 fruits). (e) Representative scanning electron micrographs showing RECs and SECs comprising the petal AZ, following manual separation at the time of petal abscission and flower PP abscission (*n* = 5). (f) Representative images of sections, DAB staining, BCECF staining, and phloroglucinol–HCl staining of the petal AZ. Samples were collected at three developmental stages: anthesis, petal abscission, and flower PP abscission. Black dashed lines and white arrowheads indicate the location of the petal AZ (*n* = 5 flowers for sections; *n* = 21 for DAB and BCECF staining; *n* = 19 for phloroglucinol– HCl staining). (g) Representative images of sections, DAB staining, and phloroglucinol–HCl staining of the PP AZ, with samples collected at three developmental stages: anthesis, petal abscission, and flower PP abscission. Black dashed lines and arrowheads indicate the location of the PP AZ (*n* = 5 pedicels for sections; *n* = 12 for DAB and phloroglucinol–HCl staining). (h) Representative scanning electron micrographs showing RECs and SECs comprising the PP AZ following manual separation during petal abscission or flower PP abscission. The white squares indicate higher-magnification regions below each panel (*n* = 5). (i) Fruit size at the time of fruit PP abscission in fruits that underwent PP abscission and those that did not. Sampling was conducted at 19–22 DPA, corresponding to the period when fruit PP abscission occurs. Each dot represents an independent fruits; data are shown as means ± SD. Statistical analysis was performed using the Mann–Whitney test (*n* = 30 or 31 fruits each). (j) Representative images of seeds (top) and seed size at the time of fruit PP abscission for fruits that underwent PP abscission and those that did not. Sampling was conducted at 19–22 DPA, corresponding to the period when fruit PP abscission occurs. Each dot represents an independent seeds; data are shown as means ± SD. Statistical analysis was performed using the Mann–Whitney test (*n* = 37 or 25 seeds each). (k) Representative images of toluidine blue staining showing the development of the protective layer in RECs of the PP AZ. Samples were collected before sealing (at fruit PP abscission) and after sealing (6 days after fruit PP abscission) (*n* = 15). (l) Representative scanning electron micrographs illustrating the development of the protective layer in RECs of the PP AZ. Samples were collected at three stages: before sealing (at fruit PP abscission), during sealing (3 days after fruit PP abscission), and after sealing (6 days after fruit PP abscission) (*n* = 5). Scale bars, 0.5 cm (c), 100 μm (e), 50 μm (sections in f, higher-magnification images in h), 500 μm (DAB, BCECF, phloroglucinol*–*HCl images in f, g, k), 200 μm (main images in h), 2 mm (j), 10 μm (l).

However, key differences emerged during the flower PP abscission stage. In *P. yedoensis*, petal abscission consistently preceded flower PP abscission, such that petals were rarely present when pedicel detachment occurred (Fig. S5). By contrast, *P. sargentii* almost always underwent flower PP abscission with the petals still attached (Fig. 5b).

Notably, petal color gradually shifted from white at anthesis to a pale reddish hue, becoming most prominent by the time of flower PP abscission (Figs. 5c, S6). Floral color change, defined as age-or pollination-dependent shifts in petal pigmentation, is a widespread strategy in angiosperms to enhance reproductive efficiency (Reverté et al., 2016, Ruxton and Schaefer, 2016, Trunschke et al., 2021). These observations suggest that successful fertilization accelerates petal abscission, whereas unfertilized flowers retain petals longer before being discarded via pedicel abscission. Consistent with this idea, petal abscission was associated with well-developed fruits, whereas flower PP abscission frequently occurred in smaller, underdeveloped fruits (Fig. 5d). The higher frequency of flower PP abscission in *P. sargentii* (71.9%) compared to *P. yedoensis* (42.6%) (Figs. 4a, 5a) supports the idea that *P. sargentii*, potentially due to lower fertilization efficiency, extends the fertilization window by delaying petal shedding.

Forcible removal of petals at the time of flower PP abscission resulted in torn and irregular surfaces, as revealed by SEM, indicating that abscission had not yet been activated (Fig. 5e). Histological analysis supported this notion: although the petal AZ was pre-formed at anthesis, we observed ROS accumulation and cellular alkalinization only when petal abscission was actively proceeding (Fig. 5f). Unlike *P. yedoensis*, which lacks lignin deposition at the petal AZ, *P. sargentii* formed a lignin brace in the SECs only during active petal abscission, suggesting that lignification is part of a post-fertilization activation program. We observed a similar pattern at the flower PP AZ, where both ROS accumulation and lignin deposition coincided with abscission onset (Fig. 5g), as confirmed by SEM (Fig. 5h).

Following fertilization, the subsequent steps appeared highly conserved between the two *Prunus* species. Calyx abscission, differences in fruit and seed development between PP-abscised and non-abscised groups, and the formation of a protective layer at the peduncle REC were all similarly observed in *P. sargentii* (Figs. 5i–l, S3).

Together, these findings demonstrate that *P. yedoensis* and *P. sargentii* both employ a hierarchical abscission program that filters reproductive structures in sequence, from petals to flower pedicels to fruit pedicels, to finely tune final fruit load. Although the overall framework is conserved, *P. sargentii* exhibits a distinct early-phase adjustment by delaying petal abscission and favoring flower PP abscission in response to fertilization variability. This species-specific modulation likely reflects an adaptive mechanism to accommodate fluctuating pollination success, enabling both species to avoid resource overinvestment and converge on a reproductively optimal fruit number (Fig. 6).

**Fig. 6.**
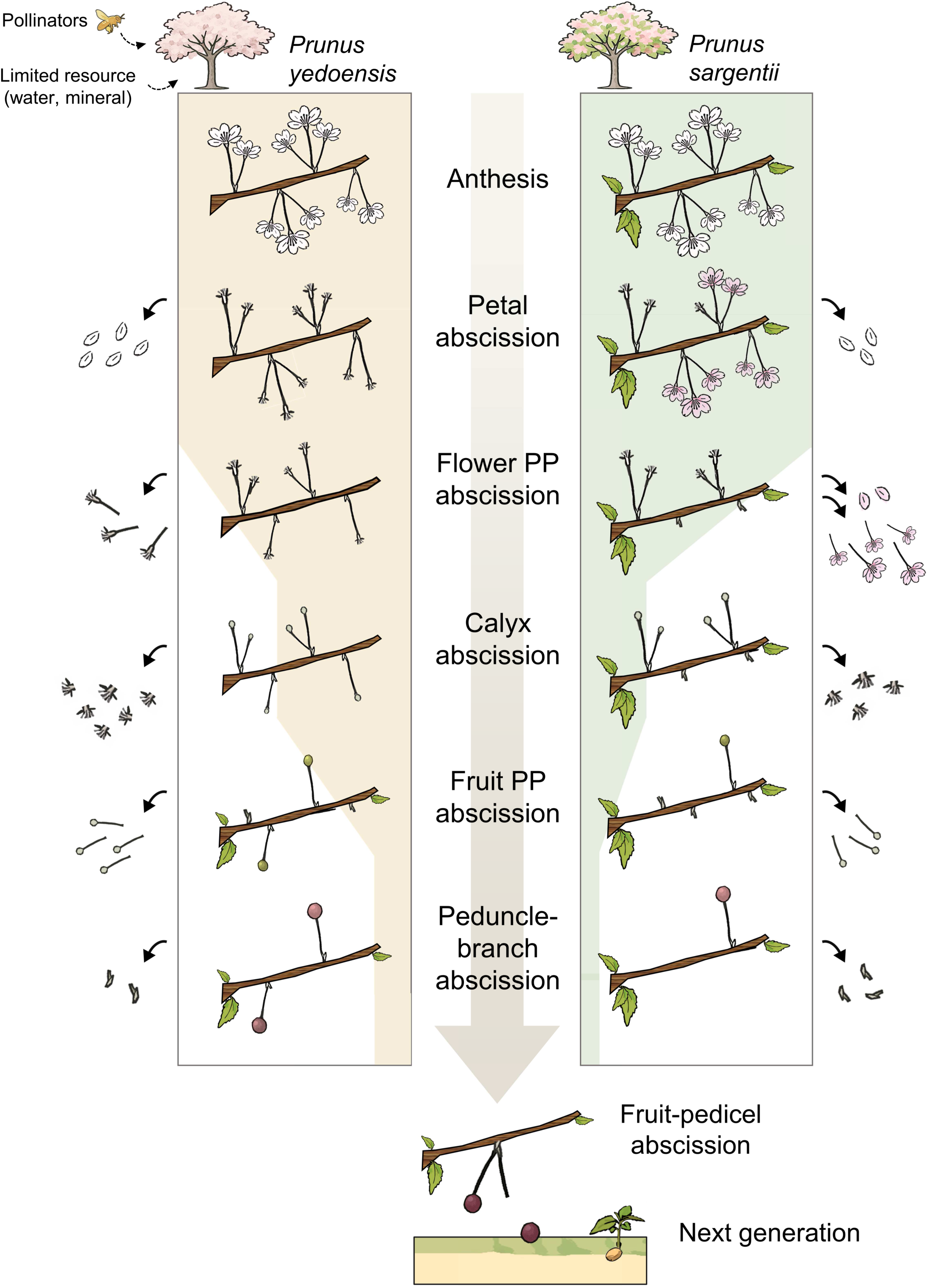
Diagram of the hierarchical abscission program regulating reproductive allocation in *Prunus yedoensis* and *Prunus sargentii*. Model illustrating five spatially and temporally distinct abscission events for petals, the calyx, flower pedicel–peduncle (PP), fruit PP, and peduncle–branches, during reproductive development in *P. yedoensis* and *P. sargentii*. Each event serves as a developmental checkpoint that filters reproductive organs based on fertilization status and resource availability. The number of flowers or fruits remaining at each stage, along with the progressively narrowing shaded background, reflects actual retention proportions. In *P. yedoensis*, petal abscission is pollination-independent, whereas petals are retained in unfertilized flowers and shed only upon fertilization in *P. sargentii*. These unfertilized flowers are removed via flower PP abscission, and even fertilized flowers undergo a second filtering step through fruit PP abscission, which eliminates a substantial proportion of developing fruits to adjust final fruit number in line with the plant reproductive capacity.

## Discussion

Among the countless flowers that bloom on a single tree, which ones are ultimately allowed to mature into fruits, and how is this decision made? This question prompted this study, inspired by the fleeting abundance of cherry blossoms in spring. Arabidopsis has been instrumental in revealing the molecular mechanisms of floral organ abscission, but it falls short when trying to capture the complexity of reproductive strategies in long-lived, cross-pollinating perennials, especially woody plants.

In this study, we investigated floral organ abscission in two *Prunus* species and identified five spatially and temporally distinct abscission events: petal, calyx, flower pedicel– peduncle (PP), fruit PP, and peduncle–branch abscissions. Rather than serving as mere developmental transitions, these events functioned as hierarchical checkpoints that sequentially filtered reproductive structures. Surplus flowers, initially produced to buffer against fertilization failure, were progressively eliminated to adjust the final fruit load in accordance with nutrient availability and pollination efficiency. This stepwise filtering program illustrates how perennial woody plants balance flower overproduction with their selective retention to align reproductive output with both environmental conditions and internal constraints. Our findings provide a conceptual framework for understanding abscission as an active strategy that modulates reproductive investment to optimize efficiency under fluctuating conditions rather than as a passive outcome of development (Fig. 6).

We observed a particularly striking divergence in the pattern of petal abscission between *P. yedoensis* and *P. sargentii*. In *P. yedoensis*, petals were shed shortly after anthesis, independently of fertilization, whereas in *P. sargentii*, petal retention was tightly linked to fertilization. This difference likely reflects broader ecological and physiological strategies*. P. yedoensis* produces a profusion of flowers that bloom before leaf emergence, possibly maximizing pollinator attraction at the cost of limited photosynthetic support. By contrast, *P. sargentii* produces flowers concurrently with leaf expansion, ensuring a more stable energy supply. Additionally, *P. sargentii* often inhabits cooler, high-altitude environments where delayed petal abscission may increase the chances of successful pollination (Chang et al., 2004).

These contrasting patterns raise a broader question: what determines the intrinsic timing of floral organ abscission? We propose that floral abscission operates under an internally governed timing program that unfolds after anthesis. This developmental trajectory likely coordinates multiple steps such as stigma senescence, ovule fate, and petal detachment, in a temporally linked manner. In this framework, fertilization does not act as the primary trigger, but rather as a modulatory input that can accelerate preexisting developmental transitions. Such a mechanism reconciles the strategies observed in different *Prunus* species: in *P. yedoensis*, the timer proceeds independently of fertilization, whereas fertilization advances timing in *P. sargentii*, ensuring that petals are only shed after reproductive success is confirmed.

This model finds support in Arabidopsis, in which floral organ abscission is delayed, but not prevented, in sterile mutants (Kandasamy et al., 2005, Kim et al., 2013, Meng et al., 2016, Furuta et al., 2024), suggesting the presence of a time-dependent mechanism. Importantly, abscission still proceeds even in mutants lacking components of the core ethylene signaling pathway or of the IDA–HAE/HSL2 module (Butenko et al., 2003, Cho et al., 2008, Stenvik et al., 2008), indicating that this temporal program operates independently of these canonical regulators. As a self-fertilizing species, Arabidopsis has drawn limited attention to fertilization-linked abscission dynamics. Yet this underexplored dimension now deserves closer investigation. Revisiting classic Arabidopsis mutants in this light may help uncover the molecular basis of a hidden layer of abscission control: one that, while playing a minor role in Arabidopsis, may be more central to shaping reproductive strategies in other species, including *Prunus*.

Another key question concerns spatial specificity: how do five temporally distinct abscission events occur in closely adjacent tissues in *Prunus*, despite the diffusible nature of signals like ethylene and small peptides? Our findings suggest that spatial precision is not governed by ethylene concentration alone, but rather by a combination of tissue-specific responsiveness (i.e., ethylene competence) and localized signaling dynamics, particularly in the PP AZ. Although the concept of ethylene competence was proposed decades ago (Osborne and Morgan, 1989), its cellular basis remained elusive. We show here that competence correlated with renewed cell division within the AZ. Although the AZ is pre-patterned by anthesis, it remains developmentally quiescent until triggered by internal cues. Upon reactivation, it undergoes cell division, followed by differentiation into RECs and SECs and, in some cases, lignin deposition, suggesting a coordinated developmental program.

Although such REC and SEC specialization was previously described in Arabidopsis (Lee et al., 2018), the upstream regulatory pathway is unknown, and it remains unclear whether renewed cell division is also required to initiate abscission in Arabidopsis. In *Prunus*, the clear onset of cell division during AZ activation raises the possibility that the AZ functions as a previously unrecognized, temporally restricted meristem. Its transient and spatially confined reactivation suggests the presence of a developmental niche, potentially governed by meristem maintenance modules akin to those controlling floral meristems. Future studies integrating transcriptomic, epigenomic, and lineage-tracing approaches will be essential to uncover how competence and plasticity are conferred to AZs and whether meristem-like regulatory logics underlie spatial specificity in abscission.

Finally, we observed that fertilization alone does not guarantee fruit retention: over 80% of all fertilized flowers were ultimately discarded. This finding indicates the presence of a stringent post-fertilization selection mechanism. In contrast to species such as Arabidopsis or *Brassica napus*, in which fruit number is largely determined during floral initiation (Walker et al., 2021), *Prunus* species can dynamically adjust their fruit number after flowering via sequential abscission events. This mechanism challenges existing notions of a fixed target fruit number and highlights the importance of post-fertilization fruit filtering. Future studies could examine how environmental or nutritional conditions alter the threshold for fruit retention, with direct implications for crop management in species like apple (*Malus domestica*) and pear (*Pyrus* sp.), where excessive fruit set often necessitates manual thinning (Costa et al., 2018). Understanding how plants determine and store decisions about fruit load could help inform more efficient agricultural management practices.

From an ecological perspective, although *Prunus* species exhibit the *r*-selected trait (suited to a reproductive strategy of rapid reproduction under high mortality) of copious flowering, the selective removal of underdeveloped fruits reflects a partial shift toward *K*-selection (slower reproduction and greater parental care in the face of competition). This potential shift aligns with the view that reproductive strategies exist along a continuum rather than as discrete categories (Pianka, 1970, Gadgil and Solbrig, 1972). By shedding low-fitness fruits and retaining fully developed, dispersal-competent fruits, *Prunus* species enhance their overall reproductive efficiency and promote effective seed dispersal (Cain et al., 2000, Nathan et al., 2008). These strategies likely evolved to balance the uncertainty of pollination with the high cost of fruit maturation in perennial systems.

Taken together, our findings suggest that organ abscission in perennial woody plant species is not merely a passive developmental conclusion, but an actively regulated program that integrates physiological status, fertilization outcomes, and ecological context. By dissecting these spatially and temporally resolved abscission programs, we begun to uncover the flexible yet finely tuned strategies that underlie reproductive success in long-lived plants. Importantly, our work highlights the critical decision points within the abscission hierarchy, especially the selective retention or elimination of fertilized flowers, as key regulatory nodes where developmental and ecological signals converge. This conceptual framework deepens our understanding of how plants optimize reproductive output and provides a strategic foundation for future studies into the molecular, cellular, and ecological controls of organ shedding in perennial systems.

## Acknowledgements

We thank Dr. Hyunmo Choi and Dr. Sung-Joon Na for advice on this work. We also thank planteditors for editorial assistance during the preparation of the manuscript. Y.L. was supported by the Suh Kyungbae Foundation (SUHF-19010003) and a National Research Foundation of Korea (NRF) grant funded by the Korean government (MSIT) (No. RS-2021-NR60084 and RS-2023-NR076399). J-ML was supported by a Hyundai Motor Chung Mong-Koo scholarship. W-TJ and J-ML were supported by the Stadelmann–Lee Scholarship Fund at Seoul National University, Korea

## Author contributions

W-TJ and YL conceived the study. W-TJ, J-AK, AC, SL, and J-ML carried out the experiments. W-TJ, J-AK, AC, SL, and JK conducted the investigation. W-TJ and AC were responsible for data visualization. YL acquired funding for the study. YL managed the project and provided supervision. W-TJ, AC, and YL drafted the original manuscript. W-TJ, J-AK, AC, SL, JK, J-ML, and YL contributed to the review and editing of the manuscript.

## Data availability

The source code for petal color acquisition, correction, and visualization is publicly available at GitHub: https://github.com/AhyeonChn/-Petal-color-acquisition-correction-visualization.

## Supporting Information

Additional Supporting Information may be found online in the Supporting Information section.

**Fig. S1.** Petal abscission is activated independently of fertilization in *Prunus yedoensis*.

**Fig. S2.** Effects of ACC treatment on petal and flower pedicel–peduncle abscission in *Prunus yedoensis*.

**Fig. S3.** Progression of calyx abscission in *Prunus* species.

**Fig. S4.** Peduncle–branch abscission in *Prunus yedoensis*.

**Fig. S5.** Flower pedicel–peduncle abscission and calyx abscission do not occur prior to petal abscission in *Prunus yedoensis*.

**Fig. S6.** Distribution of RGB values for petals at different abscission phases in *Prunus sargentii*.

**Fig. SI.**
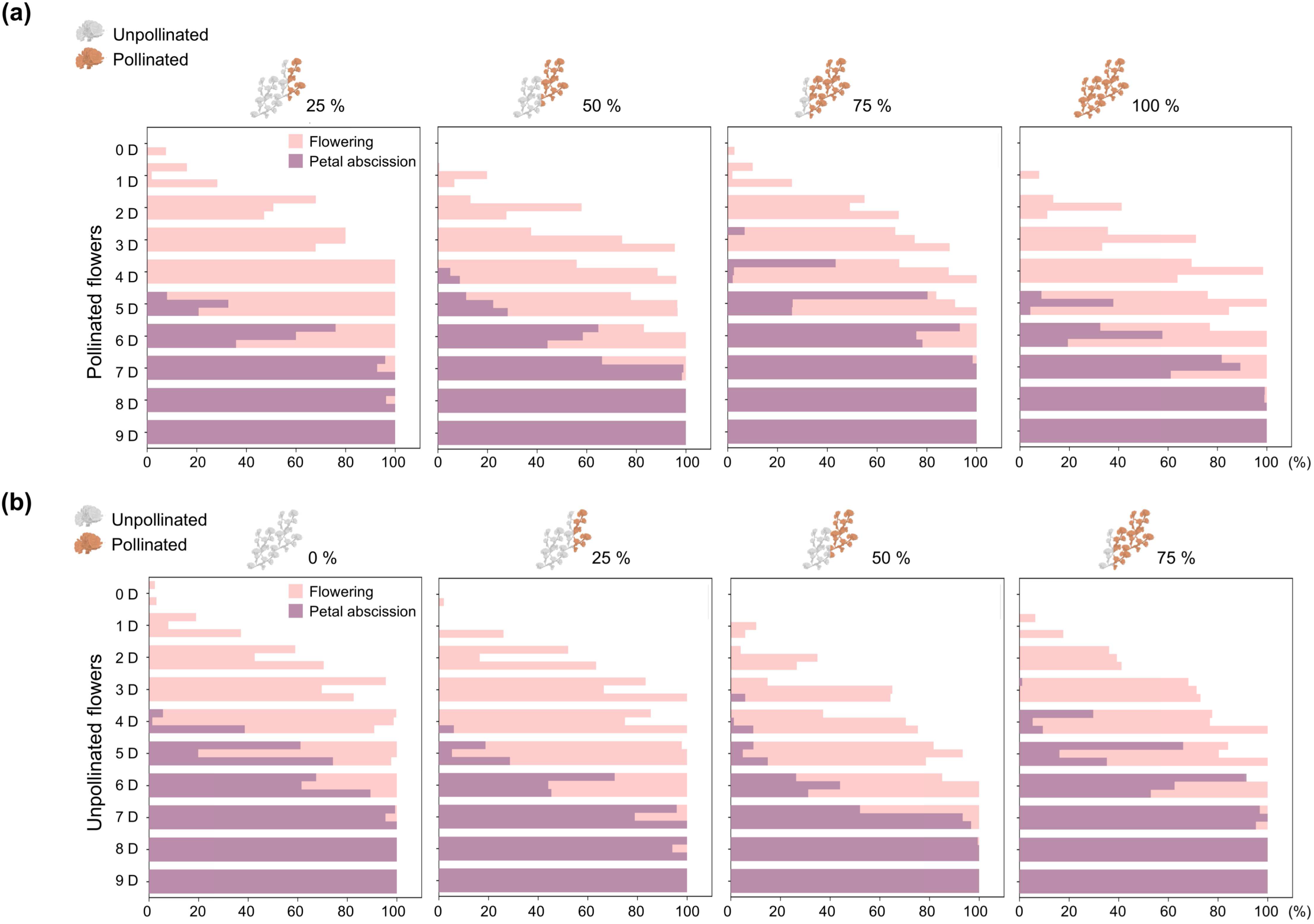
Petal abscission is activated independently of fertilization in *Prunus yedoensis*. Petal abscission kinetics were assessed under varying levels of hand pollination; results are presented separately for pollinated (a) and nonpollinated flowers (b). Cross-pollination was performed on 0%, 25%, 50%, 75%, and 100% of all flowers per tree at 0-1 day post­anthesis (DPA). Each horizontal bar represents data from an individual tree *(n* = 3 trees).

**Fig. S2.**
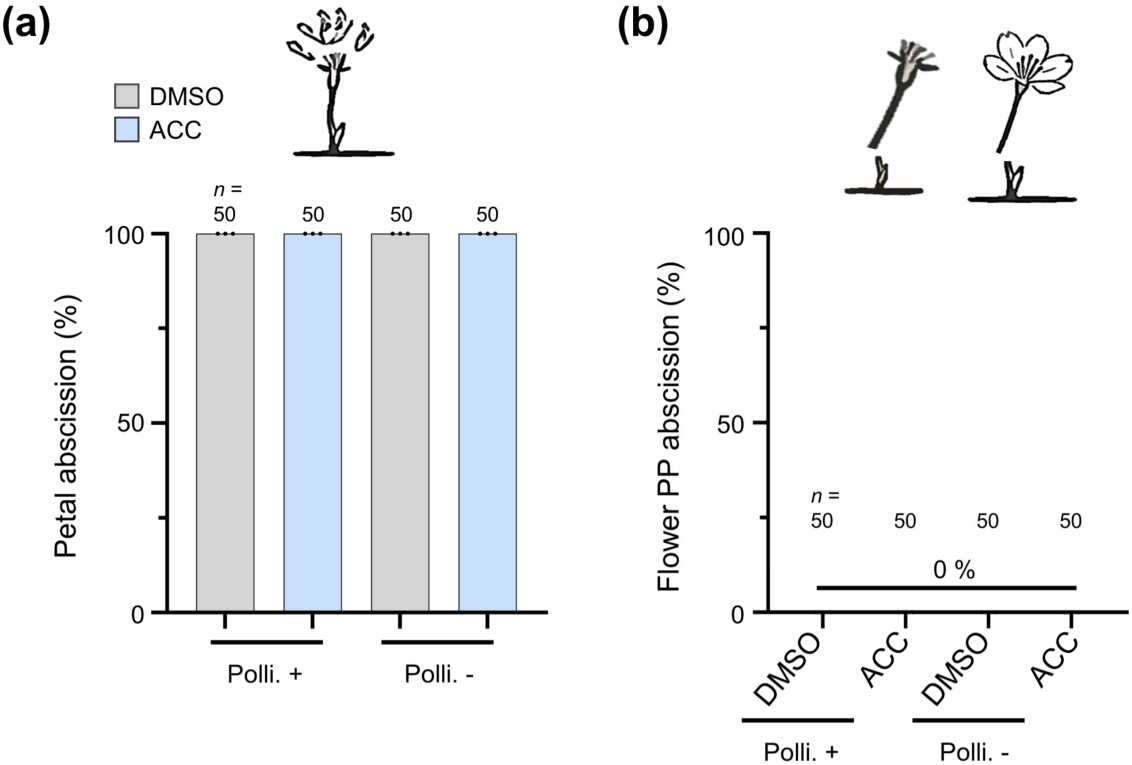
Effects of ACC treatment on petal and flower pedicel-peduncle abscission in *Prunus yedoensis*. Frequency of petal abscission (a) or flower pedicel-peduncle (PP) abscission, assessed in P. yedoensis following treatment with 100 µM 1-aminocyclopropane-1-carboxylic acid (ACC) or a mock solution (DMSO). Treatments were applied at 1 DPA, and abscission was monitored daily; abscission rates at 6 DPA are shown here, prior to the onset of natural flower PP abscission. The comparison between pollinated and nonpollinated flowers helped evaluate the influence of pollination on abscission in response to ACC treatment (« = 50 flowers and *n* = 50 flower pedicels).

**Fig. S3.**
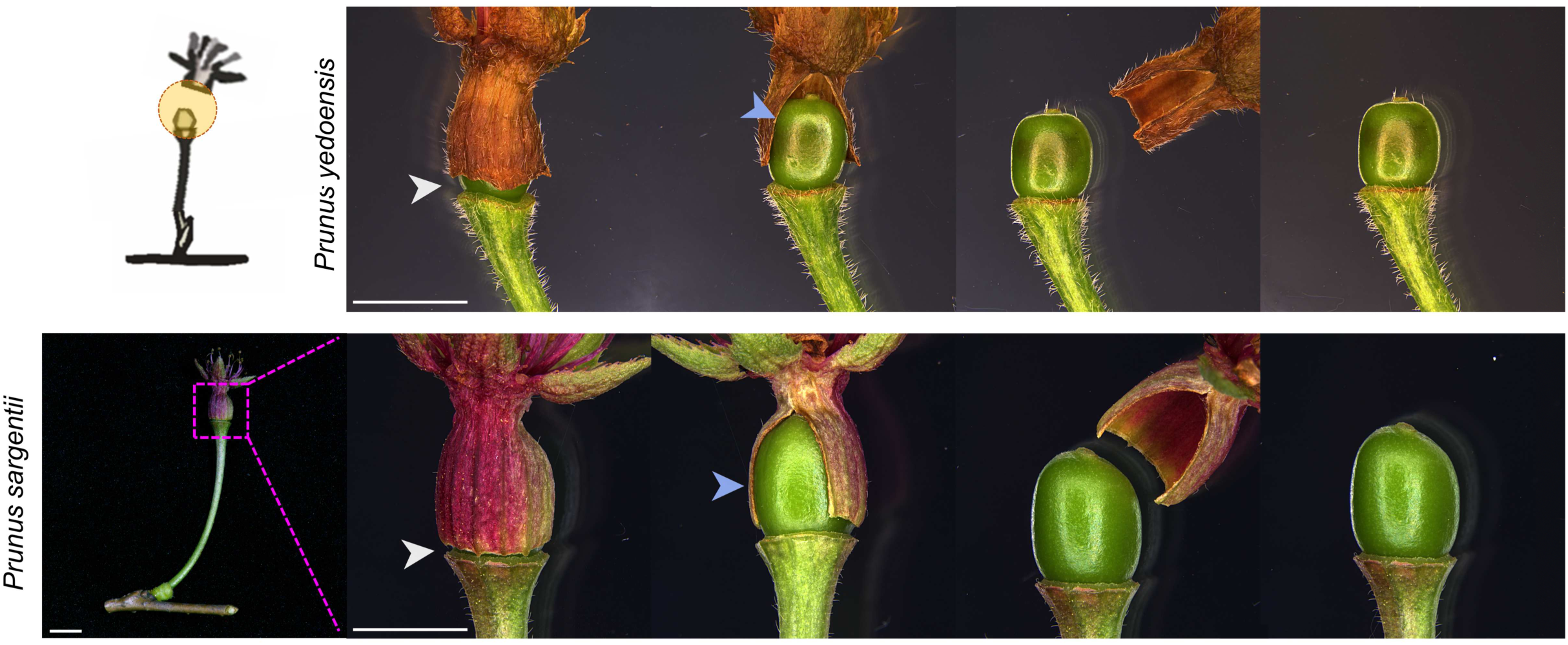
Progression of calyx abscission in *Prunus* species. Representative microscopy images illustrating calyx abscission. In Prunus species, a distinct vertical abscission line (white arrowheads) becomes visible on the calyx prior to fruit expansion, whereas a longitudinal abscission line (blue arrowheads) is not visible at this stage. As the fruit expands, the vertical line ruptures first, followed by the longitudinal line. Notably, the longitudinal line appears to form along a predetermined path, as it consistently manifests as a single cleavage site rather than as multiple lines, despite its initial lack of visibility. Ultimately, once longitudinal abscission progresses as the fruit expands, the fruit becomes exposed. The magenta box indicates the region that is shown at higher magnification to the right. Scale bars, 5 mm.

**Fig. S4.**
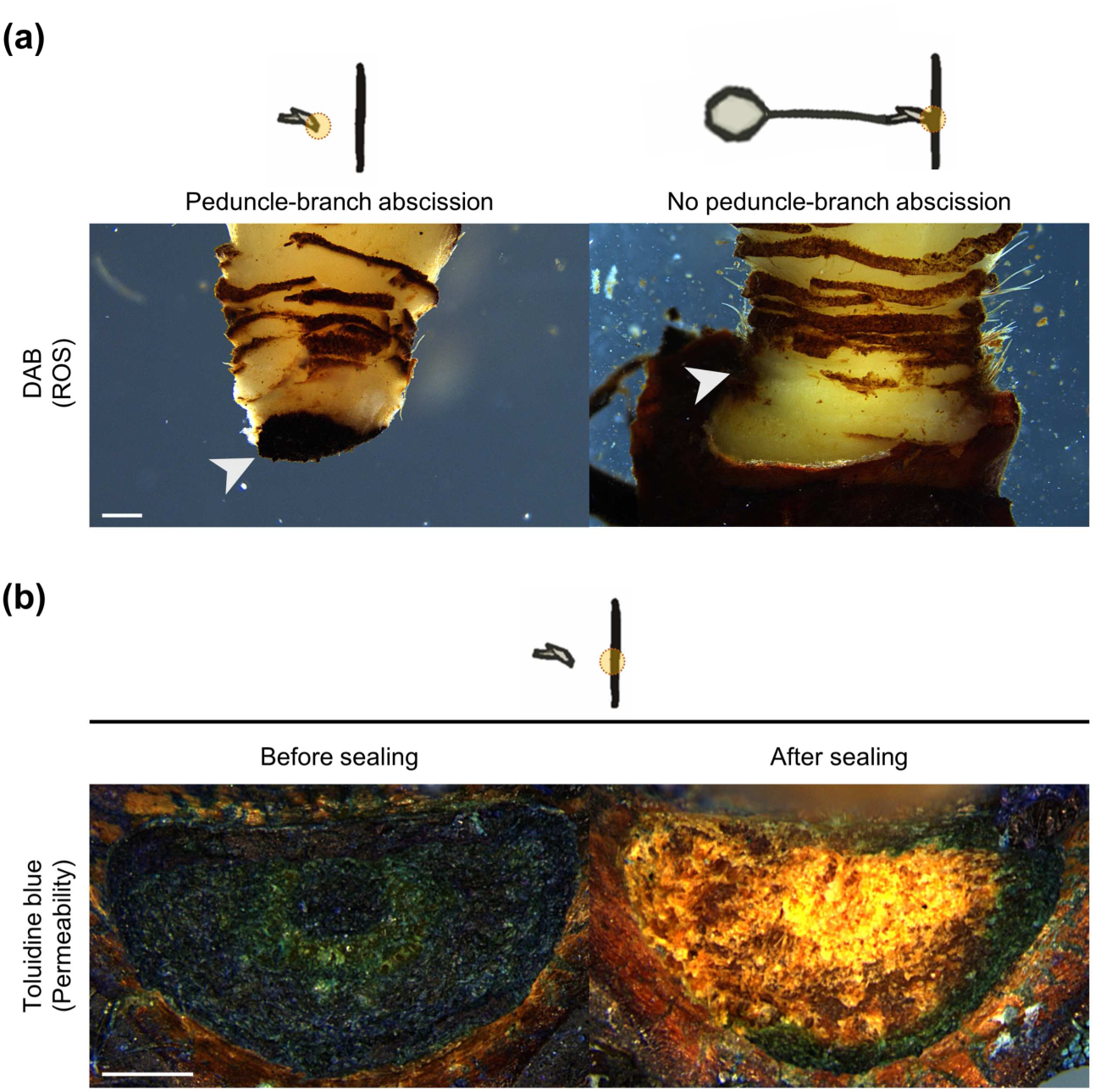
Peduncle-branch abscission in *Prunus yedoensis*. (a) DAB staining to detect ROS accumulation in secession cells at the peduncle-branch abscission zone during abscission at 30 DPA. The white arrowheads indicate the location of the abscission zone *(n -* 15). (b) Toluidine blue staining-based permeability assay to evaluate the formation of a protective layer in residuum cells immediately after peduncle-branch abscission (30 DPA; Before sealing) and at a later time point (40 DPA; After sealing) *(n* = 15). Scale bars, 500 µm.

**Fig. S5.**
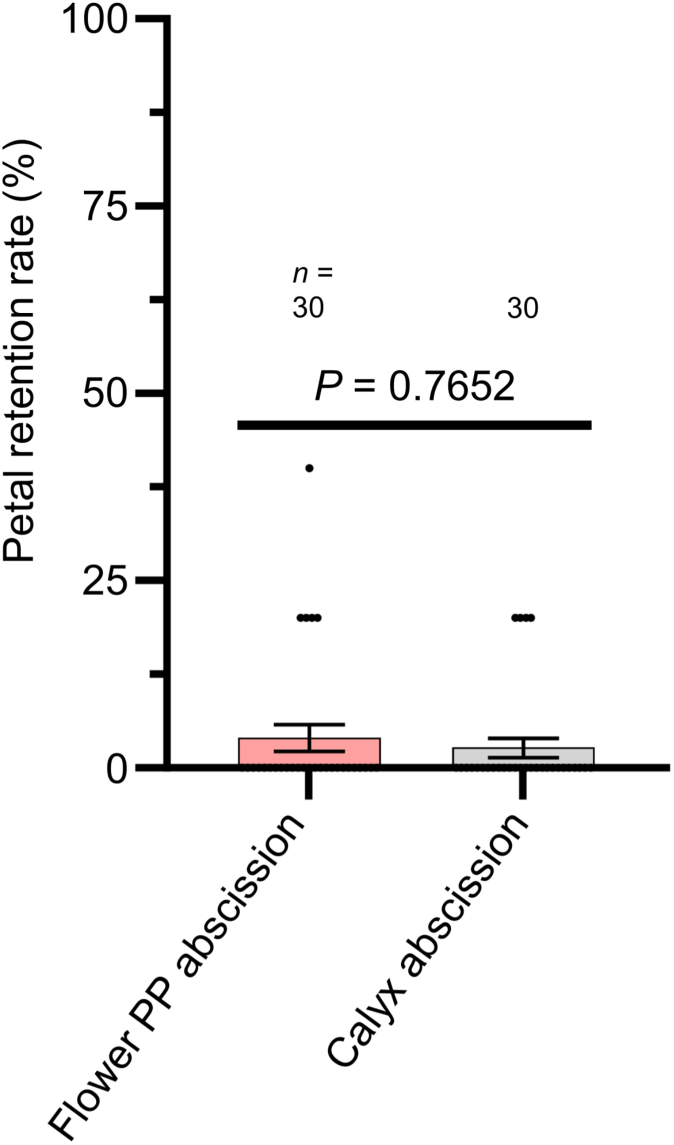
Flower pedicel-peduncle abscission and calyx abscission do not occur prior to petal abscission in *Prunus yedoensis*. The proportion of petals remaining was assessed in cases of flower PP abscission or calyx abscission *(n* = 30 flowers). A Mann-Whitney test revealed no statistically significant difference between the two groups (two-tailed P-value = 0.7652). Data are shown as means ± SD.

**Fig. S6.**
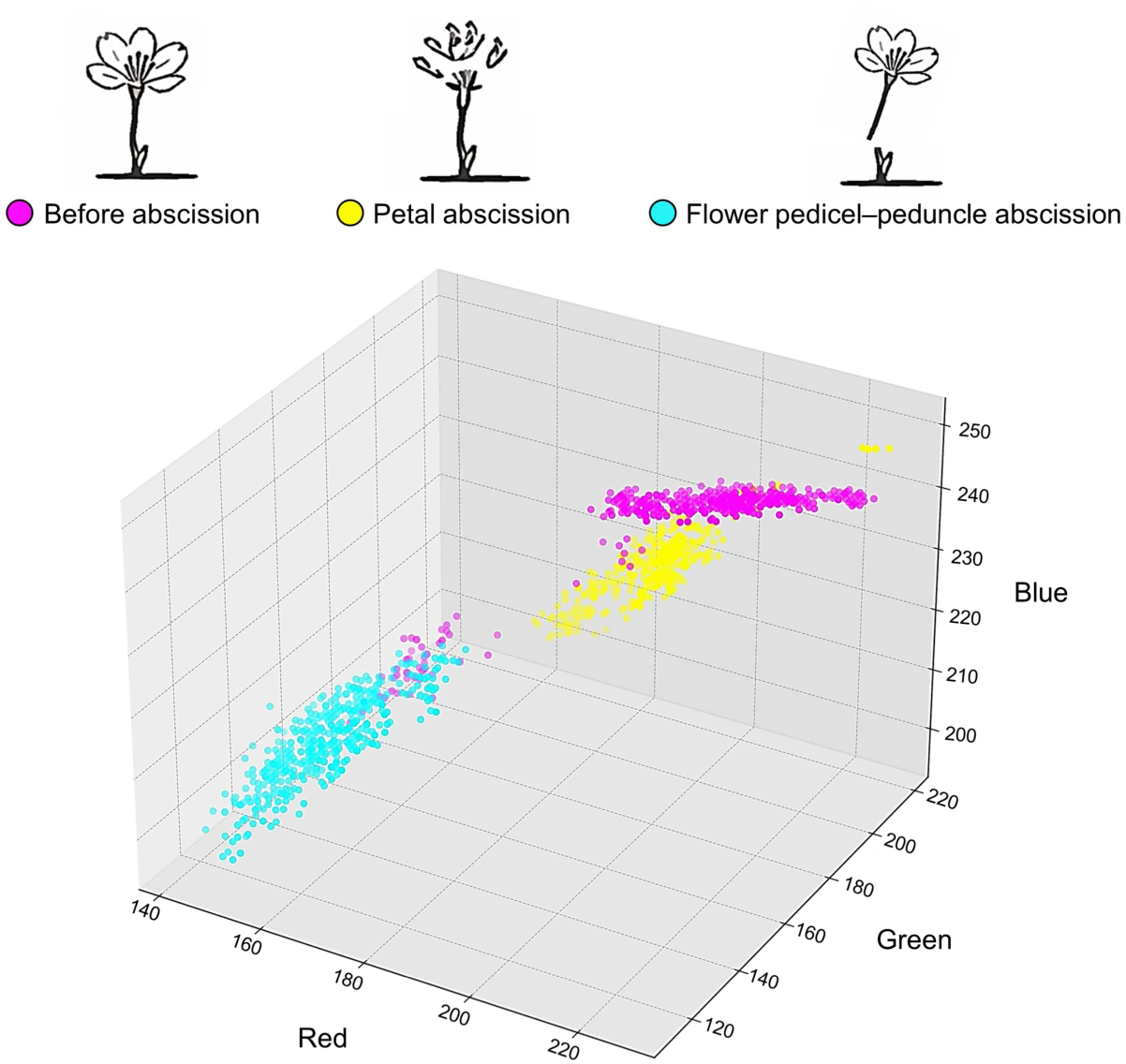
Distribution of RGB values for petals at different abscission phases in *Prunus sargentii*. 3D scatterplot showing the RGB color distribution of petals in Prunus sargentii. Each axis represents the measured red, green, and blue values. Each point corresponds to the color of one petal at a specific stage of abscission: magenta, period before abscission; yellow, petal abscission stage; cyan, after flower pedicel-peduncle abscission (w = 20 flowers).

